# Contrasting within-donor gene covariation identifies disease-associated cells and genes in case–control single-cell studies

**DOI:** 10.64898/2026.11.06.764894

**Authors:** Fengge Chang, Zhana Duren

## Abstract

In case–control single-cell studies, disease often affects only a subset of cells of a given type within each patient. Comparing all cells from patients with all cells from control donors therefore dilutes disease signals, while donor variation and recorded or unrecorded batch effects can be confounded with disease status. We present IDC (Identification of Disease-associated Cells), which uses this cellular heterogeneity as signal. A mixture of affected and unaffected cells creates coordinated variation among disease-associated genes within each patient, whereas shifts shared by all cells of a donor leave this within-donor covariance unchanged. IDC therefore contrasts within-donor gene–gene covariance between patients and controls to learn disease-associated expression programs without batch labels, then applies positive–unlabeled learning to score each cell’s disease-like state and estimate each donor’s disease burden. IDC also computes, in closed form, gene contribution scores and leave-gene-out cell scores that exclude each tested gene, enabling within-donor differential expression testing and cross-donor meta-analysis without retraining. In simulations, IDC identified affected cells and estimated donor burden more accurately than deep generative and label-refinement methods, remained robust to strong unrecorded batch effects, and detected differentially expressed genes more sensitively than case–control pseudobulk methods while controlling false discoveries. In human pancreatic islets and prefrontal cortex, disease burden in held-out donors separated type 2 diabetes (T2D) from non-diabetic donors in beta cells, and late-stage Alzheimer’s disease (AD) from non-AD donors in astrocytes and inhibitory neurons, while placing prediabetic and early-AD donors at intermediate levels. Genes with the highest IDC contribution scores were more strongly enriched for the KEGG T2D and AD pathways than case–control pseudobulk rankings, and within-donor testing revealed coordinated metabolic, cell-adhesion and synaptic changes along the inferred disease axis.

## 1 Introduction

A diagnosis is made on a person, but disease acts on cells. Within a patient’s islets or cortex, cells of the same type do not respond uniformly: some are altered by disease, while others remain close to healthy [1, 2]. Each patient therefore carries their own control. The unaffected cells share the patient’s genome, age, environment, tissue handling, and sequencing batch, and differ from their affected neighbors only in disease. Comparing the two would separate disease from nearly everything else that varies between people.

Case–control single-cell studies rarely make this comparison. Because the affected cells are unknown, analyses compare patients with healthy donors instead, typically by aggregating each donor’s cells into pseudobulk profiles [3, 4]. This choice has two costs. Unaffected cells dilute the signal from affected ones. And the comparison is made between different people, whose differences in genetics, demographics, tissue handling, and batch—many of them unrecorded—are entangled with disease. Methods that score individual cells for disease association, such as MELD [5], CNA [6], Milo [7], and HiDDEN [8], or that contrast case with control data, such as contrastive PCA [9], reduce the dilution. But they still learn what distinguishes cells of patients from those of controls, and so inherit the same confounding; moreover, testing genes between cells defined by the same expression data introduces circular inference and pseudoreplication, inflating false discoveries [10, 11, 4].

The question is whether affected cells can be found using information from within each patient. We observe that they leave a signature there. When disease alters only some cells, the mixture of altered and unaltered cells creates coordinated variation among disease-related genes within the patient, a covariance that healthy donors lack. Sample-wide shifts, such as donor background or batch, move all of that donor’s cells together and leave this within-donor covariance unchanged. The heterogeneity that dilutes conventional analyses is thus the signal, and the between-donor differences that confound them are invisible from within.

We present Identification of Disease-associated Cells (IDC), which turns this observation into a unified statistical framework (Fig. 1). First, building on contrastive covariance analysis [9], IDC contrasts donor-balanced within-donor gene–gene covariance between patients and healthy donors to extract a disease-enriched gene basis that is naturally invariant to additive donor and batch shifts. Second, projecting cells onto this basis, IDC uses positive–unlabeled (PU) learning to assign each cell a continuous disease-like score and aggregates these scores to quantify each donor’s overall disease burden. Third, because both the basis projection and the PU logistic classifier are linear transformations, IDC maps the fitted predictor directly back to gene space to compute gene contribution scores characterizing the learned disease axis and to derive closed-form leave-gene-out (LGO) cell scores without retraining the model. Using these LGO scores, IDC compares disease-like with healthy-like cells within each patient via ordered pseudobulk regression and combines slopes across donors using random-effects meta-analysis, treating donors as the units of biological replication while avoiding circular inference [10, 12].

**Figure 1.**
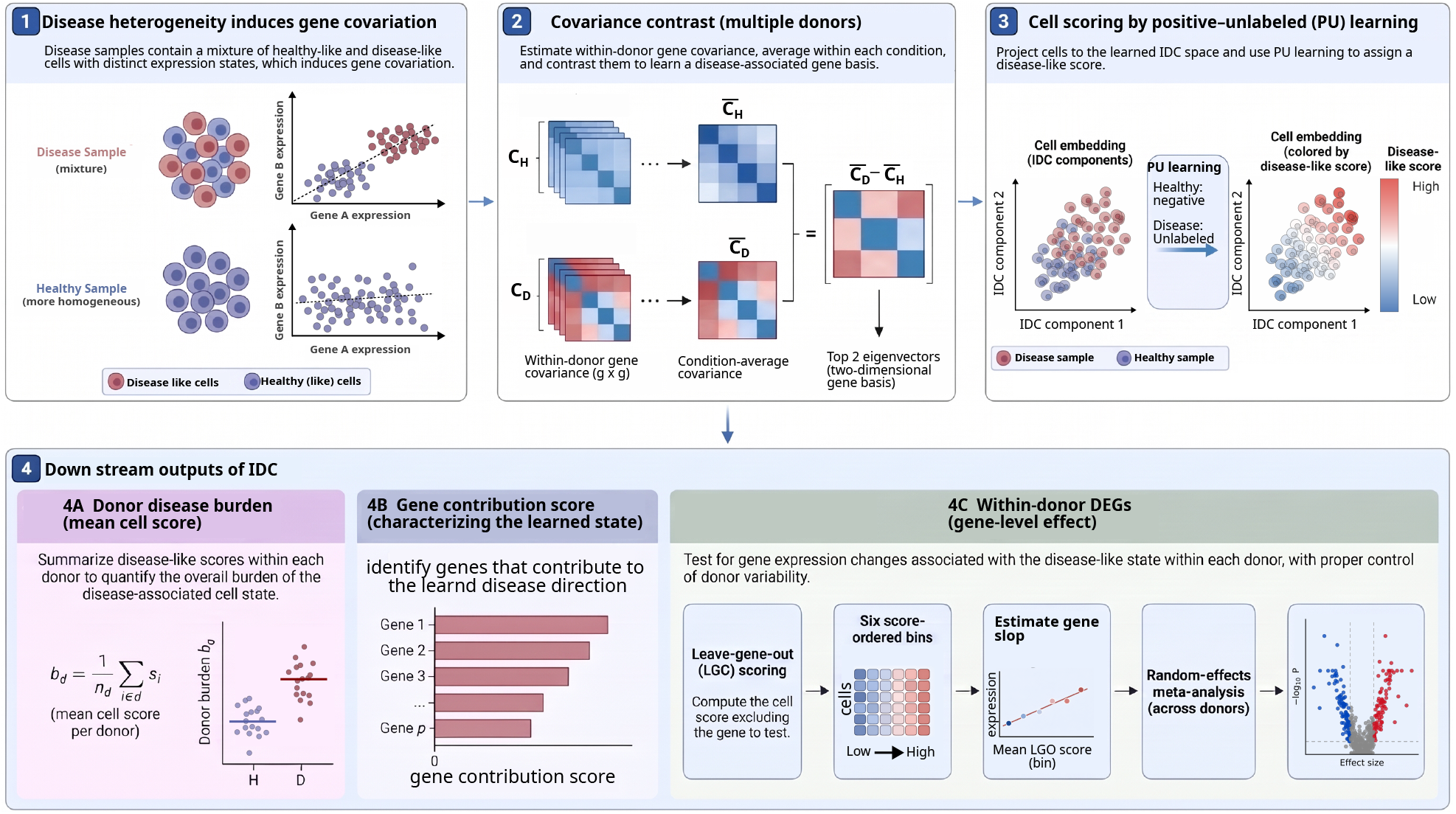
Overview of IDC. (1) Within each disease donor, mixtures of healthy-like and disease-like cells induce coordinated gene-expression variation, whereas sample-wide donor and batch shifts affect all cells of a donor together. (2) IDC computes within-donor gene–gene covariance matrices, averages them with equal donor weight within each condition, and extracts the two leading eigenvectors of the disease-minus-healthy covariance contrast. (3) Cells are projected onto this learned gene basis and assigned continuous disease-like scores by positive–unlabeled learning, treating healthy-donor cells as reference negatives and disease-donor cells as unlabeled observations. (4) Downstream outputs and gene-level inference comprise (A) donor-level disease burden, summarized by the mean cell score within each donor; and, by mapping the linear predictor back to gene space, (B) unsigned gene contribution scores characterizing the learned disease axis and (C) within-donor differential expression via closed-form leave-gene-out scoring, six score-ordered pseudobulk bins, donor-specific negative binomial regression, and random-effects meta-analysis. Leave-gene-out scoring removes the tested gene’s direct contribution in closed form without refitting the scoring model.

Together, this work makes four main contributions to single-cell case–control analysis. First, by combining contrastive within-donor covariance with PU learning, IDC resolves intra-patient cell state mixtures to score affected cells and quantify donor disease burden while automatically canceling sample-wide donor and unrecorded batch effects without requiring batch labels. Second, by pairing closed-form LGO scoring with within-donor pseudobulk regression and cross-donor meta-analysis, IDC enables each patient’s unaffected cells to serve as an internal control, roughly doubling the differential expression sensitivity of case–control pseudobulk analysis while rigorously preventing circular inference and pseudoreplication. Third, because IDC relies on lightweight spectral decomposition and evaluates all gene-specific LGO scores analytically in closed form without refitting the model, it provides a highly scalable software tool that runs an order of magnitude faster than deep generative baselines and scales readily to large multi-donor cohorts. Finally, across extensive simulations and human type 2 diabetes (T2D) pancreatic islet and Alzheimer’s disease (AD) prefrontal cortex cohorts, we show that held-out IDC donor disease burden accurately stratifies clinical disease stages—placing prediabetic and early-AD individuals between healthy and advanced disease—while IDC gene contribution scores and within-donor differential expression uncover canonical T2D and AD pathways and continuous metabolic, cell-adhesion, and synaptic transcriptional programs missed by conventional case–control comparisons.

### 2 Methods

### 2.1 Overview of the Identification of Disease-associated Cells (IDC) framework

Given case–control single-cell RNA-seq profiles of a given cell type across multiple donors, IDC aims to resolve cellular disease heterogeneity within patients and identify its underlying transcriptional drivers through three coupled stages (Fig. 1). Because disease-associated and healthy-like cells coexist within the same patient, donor diagnosis alone does not reveal individual cell states, but this mixture induces coordinated variation among disease-related genes that is absent in healthy controls.

First, to capture this signal without confounding from sample-wide donor or batch shifts, IDC computes within-donor gene–gene covariance matrices and contrasts their donor-balanced averages between disease and healthy cohorts to extract a two-dimensional disease-enriched gene basis (Fig. 1, Steps 1–2). Second, IDC projects cells onto this learned gene basis and applies positive–unlabeled (PU) learning to assign each cell a continuous disease-like score and quantify each donor’s overall disease burden (Fig. 1, Step 3 and 4A). Third, by mapping the linear predictor back to gene space, IDC derives gene contribution scores (Fig. 1, Step 4B) and closed-form leave-gene-out (LGO) cell scores, enabling within-donor differential expression testing and cross-donor meta-analysis without model retraining or circular inference (Fig. 1, Step 4C).

### 2.2 Contrastive within-donor gene–gene covariance

Within-donor covariance reflects both disease-driven state heterogeneity and baseline variation unrelated to disease, while automatically canceling sample-wide donor and batch shifts that affect all cells of a donor together. To isolate variation enriched in disease donors, IDC contrasts gene–gene covariance between disease and healthy donors (Fig. 1, Steps 1–2). We compute a covariance matrix separately for each donor and average these matrices with equal donor weight within each condition, preventing donors with more profiled cells from dominating the comparison.

Let 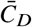 and 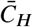 denote the average covariance matrices for disease and healthy donors, respectively. We identify gene-expression directions that maximize the difference in within-donor variance between the two conditions:

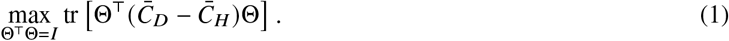

The solution is given by the two leading eigenvectors of the covariance contrast, forming an orthonormal basis Θ ∈ ℝ^*G*×2^, where *G* is the number of genes. Each eigenvector defines a data-derived gene program, with gene loadings describing the contribution of individual genes to that program. These programs capture coordinated expression variation that is more prominent within disease donors than within healthy donors, providing a low-dimensional representation of disease-enriched transcriptional variation.

### 2.3 Positive–unlabeled learning for cell scoring and donor disease burden

To represent individual cells along the learned gene programs, we standardize gene expression using donor-balanced healthy-reference means and standard deviations and project each cell onto the two eigenvectors:

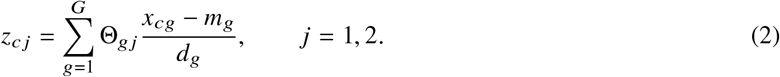

Here, *x*_*cg*_ denotes the preprocessed expression of gene *g* in cell *c, m*_*g*_ and *d*_*g*_ are its healthy-reference mean and standard deviation, and Θ_*g j*_ is the loading of gene *g* on program *j* . The resulting coordinates **z**_*c*_ = (*z*_*c*1_, *z*_*c*2_)^⊤^ define a two-dimensional representation of each cell along the disease-enriched gene programs. The healthy-reference statistics and learned basis are held fixed when projecting new cells.

The learned two-dimensional representation captures disease-enriched transcriptional variation but does not directly identify which cells are disease-like. Because disease donors may contain both disease-like and healthy-like cells, donor diagnosis alone cannot determine individual cell states. We therefore use a positive–unlabeled (PU) learning strategy, treating cells from healthy donors as reference negatives and cells from disease donors as unlabeled observations (Fig. 1, Step 3).

Because no cells are initially labeled as disease-like, we first identify candidate disease-like cells based on their Mahalanobis distance from the healthy-cell distribution in the two-dimensional embedding. These candidates provide initial positive examples for fitting a logistic regression classifier against healthy-reference cells. We then iteratively update the candidate set using the fitted classifier scores and refit the model, allowing the learned disease-like direction to extend beyond the initial distance-based selection.

The fitted classifier assigns each cell a continuous disease-like score,

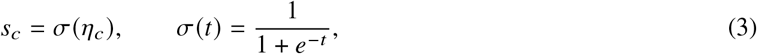

where 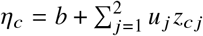 is the fitted logistic regression linear predictor with intercept *b* and program weights *u*_1_, *u*_2_ evaluated at the two-dimensional coordinates of cell *c*. Higher scores indicate greater similarity to the disease-like cellular state identified by the classifier. Finally, we quantify donor-level disease burden as the mean cell score within each donor (Fig. 1, Step 4A), which can also be evaluated on held-out donors projected onto the fixed training basis and classifier.

### 2.4 Linear gene-space decomposition and leave-gene-out within-donor differential expression

#### Gene-level decomposition and leave-gene-out scoring

Because the cell embedding is a linear projection of standardized gene expression and the classifier is linear in the resulting coordinates, the fitted predictor maps directly back to gene space. Using the effective classifier weights *u*_1_ and *u*_2_ on the two gene programs, we quantify each gene’s overall importance to the learned disease-like direction by defining a gene contribution score (Fig. 1, Step 4B) as

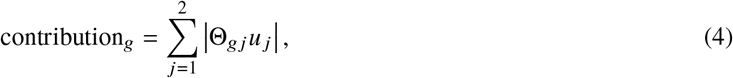

which captures the magnitude of gene *g*’s contribution across both programs without cancellation between opposing effects.

Furthermore, testing whether a gene’s expression varies with the inferred cellular disease state using scores derived from all genes introduces circularity; to eliminate this self-association, we derive a leave-gene-out (LGO) score that excludes the target gene’s direct contribution. By expanding the linear predictor as a weighted sum of individual gene expression values,

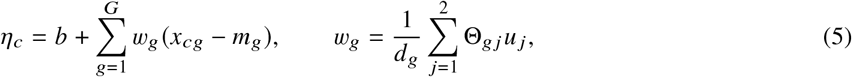

where *b* is the classifier intercept and *w*_*g*_ is the effective coefficient of gene *g* on the preprocessed expression scale, we can remove the direct contribution of any target gene *g* in closed form:

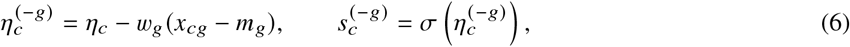

where 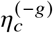 and 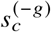 are the LGO linear predictor and disease-like score for cell *c*, respectively. Because this calculation uses the fitted model coefficients directly, LGO scores are obtained for all genes without retraining the gene programs or classifier.

#### Within-donor differential expression analysis

Using the gene-specific LGO scores, we test whether each gene’s expression varies along the inferred cellular disease continuum within each disease donor (Fig. 1, Step 4C). To mitigate single-cell count sparsity while preserving within-donor state variation, we partition cells in each disease donor *d* into six ordered bins (*k* = 1, …, 6) according to their LGO scores 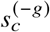. For each bin, we sum the raw counts of gene *g* across cells to construct a pseudobulk count *Y*_*gdk*_ and compute the mean LGO score *x*_*gdk*_ to represent the bin’s position along the disease-state continuum.

We then quantify the within-donor association between expression and cellular disease state by fitting a negative binomial (NB) regression separately for each disease donor *d*:

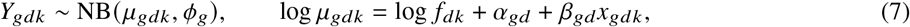

where *α*_*gd*_ is the donor-specific baseline intercept and *β*_*gd*_ is the donor-specific slope measuring how gene *g*’s expression changes toward more disease-like states. Because six bins per donor are too few to reliably estimate normalization size factors *f*_*dk*_ and gene-wise dispersions *ϕ*_*g*_ independently within each donor, we pre-estimate both parameters across all donors and hold them fixed during donor-specific fitting. Specifically, *f*_*dk*_ is computed once by the median-of-ratios method on pseudobulk bins formed from the full-model cell scores, and *ϕ*_*g*_ is estimated from a joint NB model (with donor-specific intercepts and a shared score slope) followed by empirical Bayes shrinkage toward a mean–dispersion trend across genes.

Finally, to identify genes with consistent disease-state associations across individuals while treating donors as the units of biological replication, we combine the donor-specific slope estimates 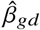 and their standard errors SE_*gd*_ across disease donors using DerSimonian–Laird random-effects meta-analysis:

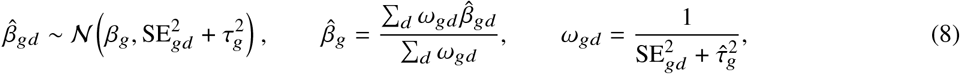

where *β*_*g*_ is the overall gene-level effect and 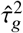 is the DerSimonian–Laird estimate of between-donor variance 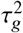 . Statistical significance of the pooled effect 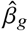 is assessed using a two-sided Wald test, followed by Benjamini–Hochberg FDR correction across genes.

### 2.5 Simulation and comparators

To benchmark IDC against known ground truth at the cell, donor, and gene levels, we used Splatter [13] to simulate 5,000 genes across 30 healthy and 30 disease donors based on parameters estimated from ROSMAP non-AD nuclei. Healthy donors contained only unaffected Group 1 cells, whereas disease donors contained mixtures of unaffected Group 1 and affected Group 2 cells, with each disease donor’s affected-cell fraction sampled independently from Uniform 0.1, 0.9 . A subset of genes were simulated as true differentially expressed genes (DEGs) between unaffected (Group 1) and affected (Group 2) cells. To systematically vary the magnitude of expression differences in these true DEGs alongside unrecorded donor-level confounding, we crossed four disease-effect strengths (`de.facLoc` ∈ {0.1, 0.3, 0.5, 0.7}, controlling the fold-change magnitude of true DEGs between Groups 1 and 2) with nine donor-specific batch-effect strengths (`batch.facLoc` ∈ {0.1, …, 0.9}, simulating unrecorded sample-level covariates by treating each donor as a separate batch), yielding 36 simulation settings. In addition to analyzing the full cohort (30 donors per condition) across all 36 settings, we evaluated the effect of sample size in the representative setting (de.facLoc = 0.5, batch.facLoc = 0.5) by subsampling 5, 10, 15, 20, or 25 donors per condition (100 random subsets per size for cell-level evaluation and 5 subsets per size for gene-level evaluation).

We evaluated performance across three levels. At the cell level, we measured the discrimination between affected and unaffected cells within disease donors using the area under the receiver operating characteristic curve (AUROC) and the area under the precision–recall curve (AUPRC). At the donor level, we assessed whether estimated donor disease burden (mean cell score) tracked the true fraction of affected cells via Pearson correlation. At the gene level, we evaluated differential expression recovery using AUPRC against the ground-truth differentially expressed genes (DEGs) and assessed FDR control via the observed false discovery proportion (FDP)—the fraction of false positives among genes called significant at a Benjamini–Hochberg adjusted *P* < 0.05 threshold—as well as observed FDP across varying nominal FDR thresholds.

To test generalization to unseen individuals, we evaluated both in-sample models (fitted and evaluated on the same cohort) and held-out donor models using five-fold donor-disjoint cross-validation. For held-out gene-level evaluation, out-of-fold IDC+PU parameters were used to compute gene-specific LGO scores on unseen disease donors, followed by within-donor negative binomial regression and random-effects meta-analysis.

For affected-cell identification, we compared IDC against scVI [14], scANVI [15], and HiDDEN [8]. To isolate the effect of the embedding representation, we applied the exact same PU learning procedure to scVI and scANVI latent spaces (‘scVI+PU’ and ‘scANVI+PU’), while evaluating HiDDEN using its native label-refinement pipeline. For gene recovery, we compared IDC’s within-donor test against conventional case–control differential expression methods—DESeq2 [16], edgeR [17], and limma-voom [18]—applied to donor-level pseudobulk counts between disease and healthy cohorts, as well as Wilcoxon rank-sum tests on HiDDEN-refined cell labels. Computational runtime was benchmarked on the representation-learning step across methods.

### 2.6 Real-data analysis and enrichment

We applied IDC to two human single-nucleus cohorts spanning healthy controls, intermediate stages, and diagnosed disease: (i) pancreatic islets from six non-diabetic (Non), eight prediabetic (Pre), and six type 2 diabetes (T2D) donors (GSE200044) [19], fitted separately per endocrine cell type on Non and T2D donors; and (ii) ROSMAP prefrontal cortex (PFC) and entorhinal cortex (EC) nuclei across seven major cell classes from non-AD, early-AD, and late-AD individuals [20], fitted separately per region and cell class on non-AD and late-AD donors with at least ten nuclei. For model fitting, we retained GENCODE v44 protein-coding genes with mean donor-pseudobulk counts per million greater than one among training donors, excluding mitochondrially encoded and cytosolic RPL/RPS ribosomal-protein genes (which remained in the downstream differential expression testing universe).

To evaluate donor-level generalization without data leakage, we used leave-one-donor-out (LODO) cross-validation across training groups (Non vs. T2D and non-AD vs. late-AD) and tested diagnostic discrimination on held-out donor burden scores using a one-sided exact Mann–Whitney test. Intermediate-stage donors (Pre and early-AD) were excluded from training and scored by averaging predictions across all frozen LODO models for the corresponding cell type. To assess biological relevance of the learned disease axis, we performed one-sided hypergeometric over-representation analysis on the top 500 genes ranked by IDC gene contribution score against all ranked genes as background, comparing fold enrichment against the top 500 genes ranked by absolute test statistics from case–control pseudobulk DESeq2 (Wald statistic), edgeR 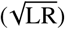, and limma-voom (moderated *t* statistic).

## 3 Results

### 3.1 IDC identifies affected cells and donor burden in simulation

We first evaluated whether IDC could resolve affected cells within disease donors containing mixtures of affected and unaffected cells. In the representative simulation setting (disease-effect and batch-effect strengths both 0.5), the two-dimensional IDC representation cleanly separated true cellular states, and the learned disease-like scores distinguished affected from unaffected cells (Fig. 2A,B). Quantitatively, IDC disease-like score achieved an AUROC of 0.956 and an AUPRC of 0.960, outperforming scVI+PU (0.869/0.863), scANVI+PU (0.657/0.653), and HiDDEN (0.784/0.793) evaluation (Fig. 2C,D). Because scVI+PU and scANVI+PU use the exact same PU classifier as IDC, these gains demonstrate that contrasting within-donor gene covariance isolates disease-associated state variation much more effectively than general-purpose deep generative embeddings. Moreover, computing the IDC representation was 10–37 times faster than fitting scVI on the same GPU across datasets of 50,000 to 250,000 cells (Fig. <u>S1</u>).

**Figure 2.**
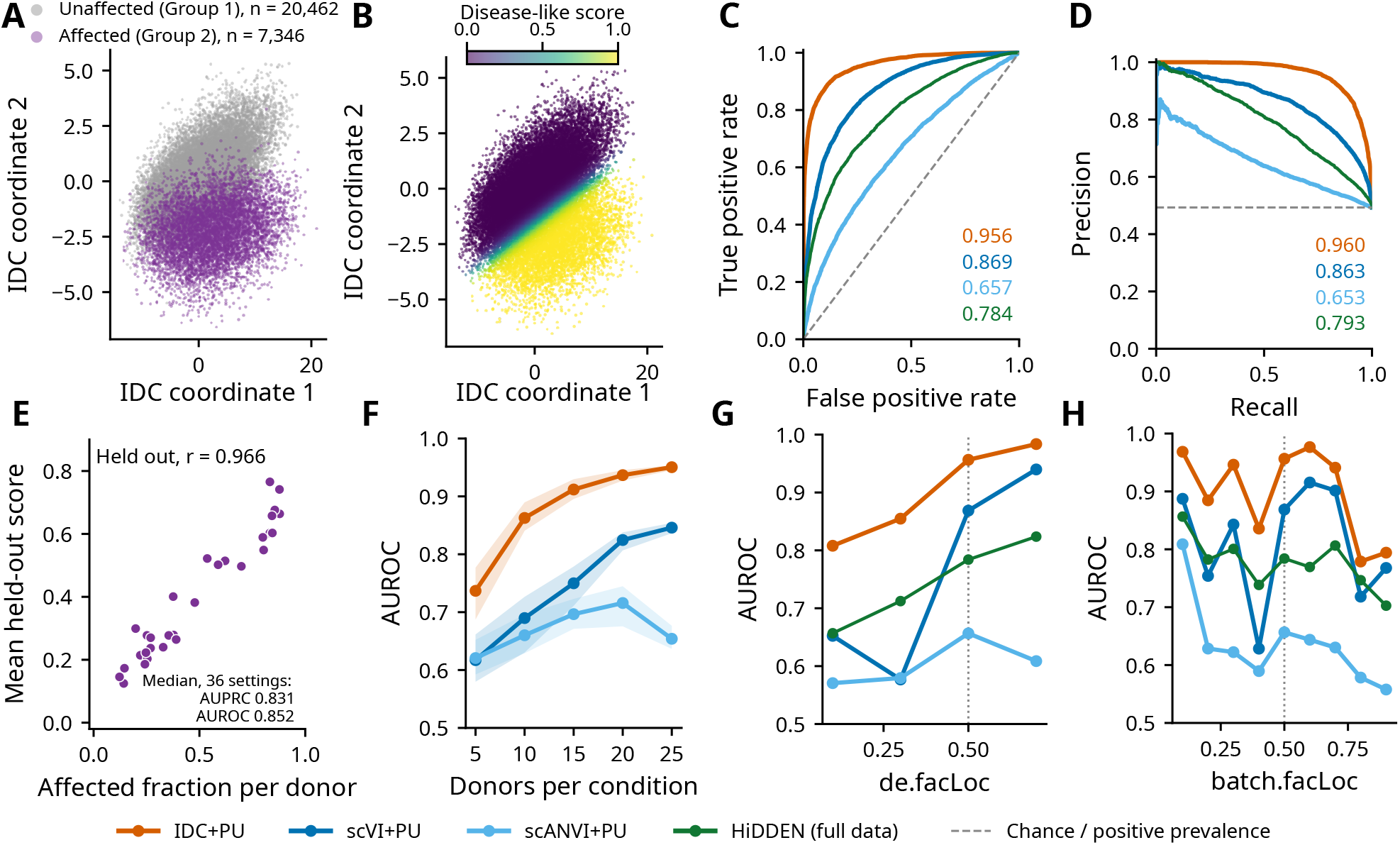
IDC identifies affected cells and estimates donor burden in simulation. (A,B) IDC coordinates in the representative setting (30 healthy and 30 disease donors; disease-effect and batch-effect strengths both 0.5), colored by true cell state (A) or disease-like score (B). (C,D) Receiver operating characteristic (C) and precision–recall (D) curves for identifying affected cells among cells from disease donors; annotations give AUROC and AUPRC, respectively. Dashed lines indicate chance discrimination or affected-cell prevalence. (E) Mean held-out disease-like score versus true affected-cell fraction for disease donors, using five donor-disjoint folds; Pearson *r* = 0.966. Inset values summarize held-out cell-identification performance across the 36 settings, evaluated within disease donors. (F) AUROC versus donors per condition in the representative setting, showing the median and interquartile range over 100 subsets per donor count. (G,H) AUROC versus disease-effect strength at batch strength 0.5 (G) and batch-effect strength at disease effect 0.5 (H) scVI and scANVI embeddings use the same PU procedure as IDC.

We next asked whether IDC could generalize to unseen individuals to estimate patient-level disease burden. Using five-fold donor-disjoint cross-validation, the mean held-out cell score per disease donor closely tracked the true affected-cell fraction in the representative setting (Pearson *r* = 0.966; Fig. 2E), while maintaining strong held-out cell discrimination across all 36 simulation settings (median AUROC 0.852, median AUPRC 0.831)(Fig. <u>S2</u>).

Finally, we evaluated robustness across challenging regimes by varying cohort sample size, disease-effect strength, and unrecorded donor-specific batch effects (Fig. 2F–H, Figs. <u>S2</u>, <u>S3</u>). Across subsampled cohorts, IDC maintained a consistent lead of 0.104–0.173 in median AUROC over the best competing method (scVI+PU), rising from 0.737 vs. 0.617 with only five donors per condition to 0.950 vs. 0.846 with 25 donors per condition (Fig. 2F). When varying disease-effect strength at fixed batch strength (0.5), IDC rose sharply even at modest disease signal (de.facLoc = 0.3), where scVI+PU, scANVI+PU, and HiDDEN remained near chance levels (Fig. 2G). Most notably, under increasing donor-specific batch effects at fixed disease effect (0.5), competing methods degraded substantially as unrecorded sample-level confounding grew, whereas IDC remained highly stable—retaining AUROCs of 0.779 and 0.794 even at the strongest batch strengths of 0.8 and 0.9, respectively (Fig. 2H)—because sample-wide shifts cancel within each donor’s covariance calculation without requiring explicit batch correction.

### 3.2 Within-donor testing improves gene recovery with calibrated FDR control

Accurate identification of affected cells does not automatically guarantee reliable recovery of the underlying differ-entially expressed genes (DEGs). We therefore benchmarked IDC’s leave-gene-out (LGO) within-donor differential expression test against conventional donor-level case–control pseudobulk methods (DESeq2, edgeR, and limma-voom) and Wilcoxon rank-sum testing on HiDDEN-refined cell labels. Across both varying disease-effect strengths (Figs. 3A, <u>S4</u>) and unknown donor-specific batch-effect strengths (Figs. 3B, <u>S4</u>), IDC roughly doubled the gene-recovery AUPRC of all competing methods. Whereas donor-level case–control pseudobulks dilute the disease signal by pooling unaffected and affected cells together—reaching AUPRCs of only 0.18–0.44 across disease-effect strengths—IDC directly contrasts bins along the inferred within-donor disease continuum, increasing AUPRC from 0.41 at weak disease signal (`de.facLoc` = 0.1) to 0.77 at strong signal (`de.facLoc` = 0.7; Fig. 3A) and maintaining a substantial lead across batch-effect strengths up to 0.7 before moderating at extreme batch levels (0.8–0.9; Fig. 3B).

**Figure 3.**
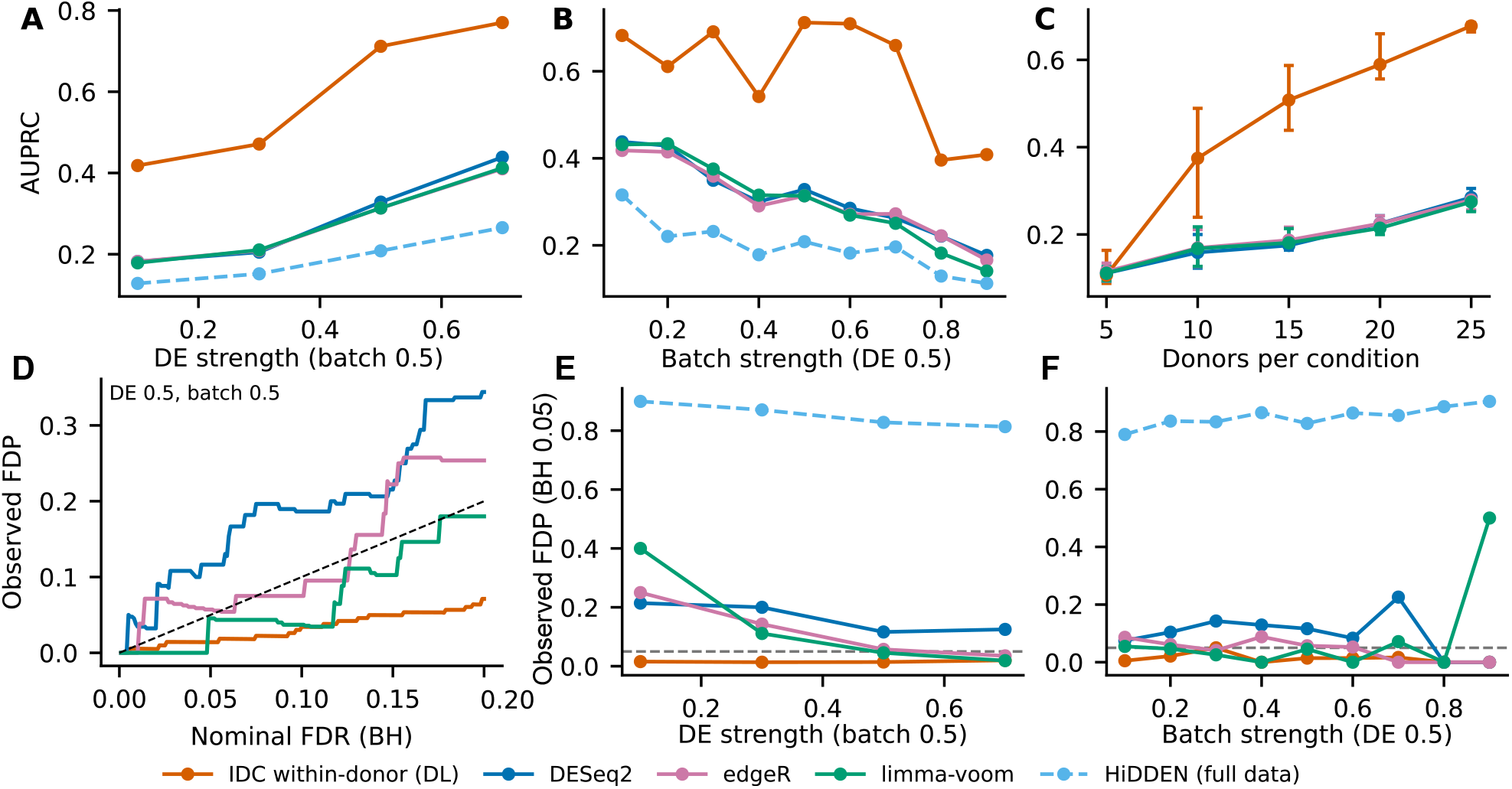
Gene recovery and observed false discovery proportion in simulation. (A,B) Gene-recovery AUPRC versus disease-effect strength at batch strength 0.5 (A) and batch-effect strength at disease effect 0.5 (B). (C) AUPRC versus donors per condition in the representative setting (disease effect 0.5, batch effect 0.5); points show means and error bars the minimum–maximum over five donor subsets. (D) Observed false discovery proportion (FDP) versus the nominal Benjamini–Hochberg threshold in the representative setting; the diagonal indicates equality. (E,F) Observed FDP at BH 0.05 versus disease-effect (E) and batch-effect (F) strength, with the other parameter fixed at 0.5; horizontal dashed lines mark 0.05. IDC uses the within-donor DerSimonian–Laird test, whereas DESeq2, edgeR and limma-voom compare donor pseudobulks between conditions. HiDDEN results in A,B,E,F use Wilcoxon tests between cell labels. FDP is FP/(TP + FP) conditional on at least one discovery; no-discovery results are undefined, not zero.

Because IDC treats donors as the units of biological replication via random-effects meta-analysis, its power scales strongly with cohort size (Fig. 3C). With five donors per condition, IDC achieved a mean AUPRC of 0.107, comparable to 0.111–0.115 for the case–control methods. However, at ten donors per condition, IDC’s AUPRC jumped to 0.375 compared with 0.159–0.169 for the alternatives, and continued to climb steadily with sample size—reaching 0.68 at 25 donors per condition while DESeq2, edgeR, and limma-voom remained below 0.29. These results show that while case–control pseudobulk methods remain bottlenecked by cell-state dilution even as cohort size grows, IDC’s within-donor test translates additional donor replication into more than a twofold gain in gene-recovery accuracy.

Crucially, this gain in sensitivity did not come at the expense of false discovery rate (FDR) control. In the representative setting, IDC’s observed false discovery proportion (FDP) remained well below nominal Benjamini– Hochberg FDR thresholds across the entire curve (0.014 at nominal FDR 0.05 and 0.034 at 0.10; Fig. 3D). Across varying disease-effect and batch-effect regimes at nominal 5% FDR, IDC consistently controlled observed FDP below 0.05 (median 0.020 across settings; Fig. 3E,F, Fig. <u>S5</u>). By contrast, conventional case–control pseudobulk methods exhibited substantial FDR inflation (observed FDP reaching 0.15–0.50) due to unrecorded donor-level batch confounding, and HiDDEN’s cell-level Wilcoxon test suffered from severe false discovery inflation (observed FDP > 0.80 across all conditions) caused by post-selection double-dipping and cell-level pseudoreplication (Fig. 3E–F). Together, these results demonstrate that combining closed-form LGO scoring with within-donor negative binomial regression and donor-level meta-analysis achieves both high power and well-calibrated FDR control.

### 3.3 Type 2 diabetes: a beta-cell state and its within-donor program

We first examined whether IDC could detect disease-associated cellular signals in human pancreatic islets across four endocrine cell types. Using leave-one-donor-out scores, we compared donor-level disease burden between non-diabetic and T2D individuals. Beta cells showed a significant increase disease burden in T2D donors (one-sided Mann–Whitney *P* = 0.0043), whereas alpha, delta, and gamma cells showed no significant differences (Fig. 4A). Prediabetic donors, excluded from model fitting, were also projected to examine their scores along the disease spectrum. We next examined how the T2D-associated signal was distributed among individual beta cells. Although cells from non-diabetic, prediabetic, and T2D donors largely overlapped in the two-dimensional representation (Fig. 4B), cells with high disease-like scores originated predominantly from prediabetic and T2D donors (Fig. 4C). Consistently, the distribution of the PU linear predictor was shifted toward higher values in T2D cells relative to non-diabetic cells, despite substantial overlap between groups (Fig. 4D). These observations indicate that the T2D-associated signal reflects a shift in the distribution of cellular states rather than a discrete separation by diagnosis.

**Figure 4.**
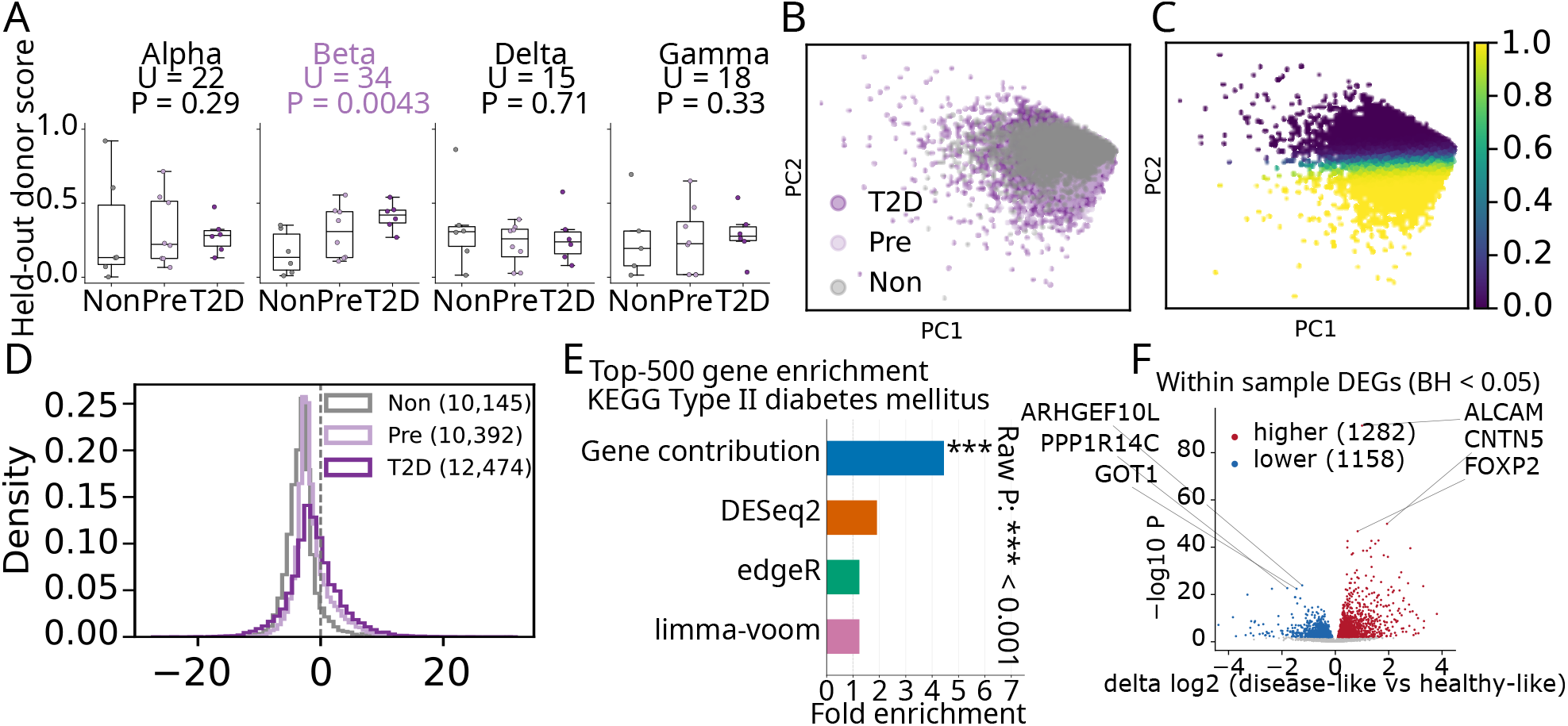
Disease-associated beta-cell states in human pancreatic islets. (A) Donor scores for four endocrine cell types, ordered Non, Pre and T2D. Non and T2D scores use leave-one-donor-out evaluation; prediabetic donors were excluded from fitting and their scores averaged across frozen LODO models. Points represent donors; boxes show medians and interquartile ranges, with whiskers extending to the most extreme observations within 1.5 interquartile ranges. *U* and raw *P* values are from exact one-sided Mann–Whitney tests of T2D > Non; Pre scores are descriptive. (B,C) Principal-component views of the IDC embedding of beta cells, colored by diagnostic group (B) or disease-like score (C). (D) Distributions of the PU logistic linear predictor by group; legend counts give numbers of cells. Pre cells in B–D were projected without fitting. (E) Over-representation of KEGG Type II diabetes mellitus among the top 500 genes ranked by the IDC gene contribution score, absolute DESeq2 Wald statistic, square root of the edgeR likelihood-ratio statistic, or absolute limma-voom moderated *t* statistic. Fold enrichment is the pathway fraction among the top 500 divided by its fraction among all genes ranked by that method; backgrounds are method-specific. Asterisks denote raw hypergeometric *P* < 0.001 (***); unmarked bars do not meet this threshold. This unsigned ranking analysis is distinct from signed GSEA and within-donor differential expression. (F) Within-donor beta-cell volcano plot: reported Δ − log_2_ effect-size summary versus log_10_ *P*. Red and blue indicate genes higher (1,282) or lower (1,158) in disease-like cells at BH < 0.05; gray indicates other tested genes. Selected labeled genes are shown for readability.

To investigate the biological relevance of the inferred disease-like state, we examined whether genes contributing to the learned direction were enriched for established diabetes-related pathways. Among the top 500 genes ranked by IDC gene contribution score, the KEGG Type II diabetes mellitus pathway showed strong enrichment (fold enrichment 4.45; hypergeometric *P* < 0.001; (Fig. 4E, Fig. <u>S6</u>), exceeding that obtained from the corresponding top-ranked genes of DESeq2, edgeR, and limma-voom. This finding suggests that the learned disease-like direction captures transcriptional features related to established T2D biology.

We next examined which genes were differentially expressed along the inferred disease-like state using IDC’s within-donor test. Among the most significant genes in pancreatic beta cells, GOT1 showed lower expression in disease-like cells, whereas ALCAM showed higher expression (Fig. 4F). GOT1 encodes an enzyme involved in the malate–aspartate shuttle and has been implicated in insulin secretion in experimental beta-cell models, while ALCAM encodes a cell-adhesion molecule. These findings reveal transcriptional changes associated with beta-cell metabolism and cell adhesion along the inferred disease-like state.

### 3.4 Alzheimer’s disease: an inhibitory-neuron state

We next applied IDC to single-nucleus RNA-seq data from the prefrontal cortex of individuals with Alzheimer’s disease (AD). To identify cell classes exhibiting disease-associated signals, we compared leave-one-donor-out disease-like scores between non-AD and late-AD donors across seven major cell classes. Astrocytes and inhibitory neurons showed significantly higher scores in late-AD donors (one-sided Mann–Whitney *P* = 0.0037 and *P* = 0.00053, respectively), whereas the remaining five classes showed no significant differences (Fig. 5A). Early-AD donors, excluded from model fitting, were projected to examine disease-state scores at an intermediate disease stage.

**Figure 5.**
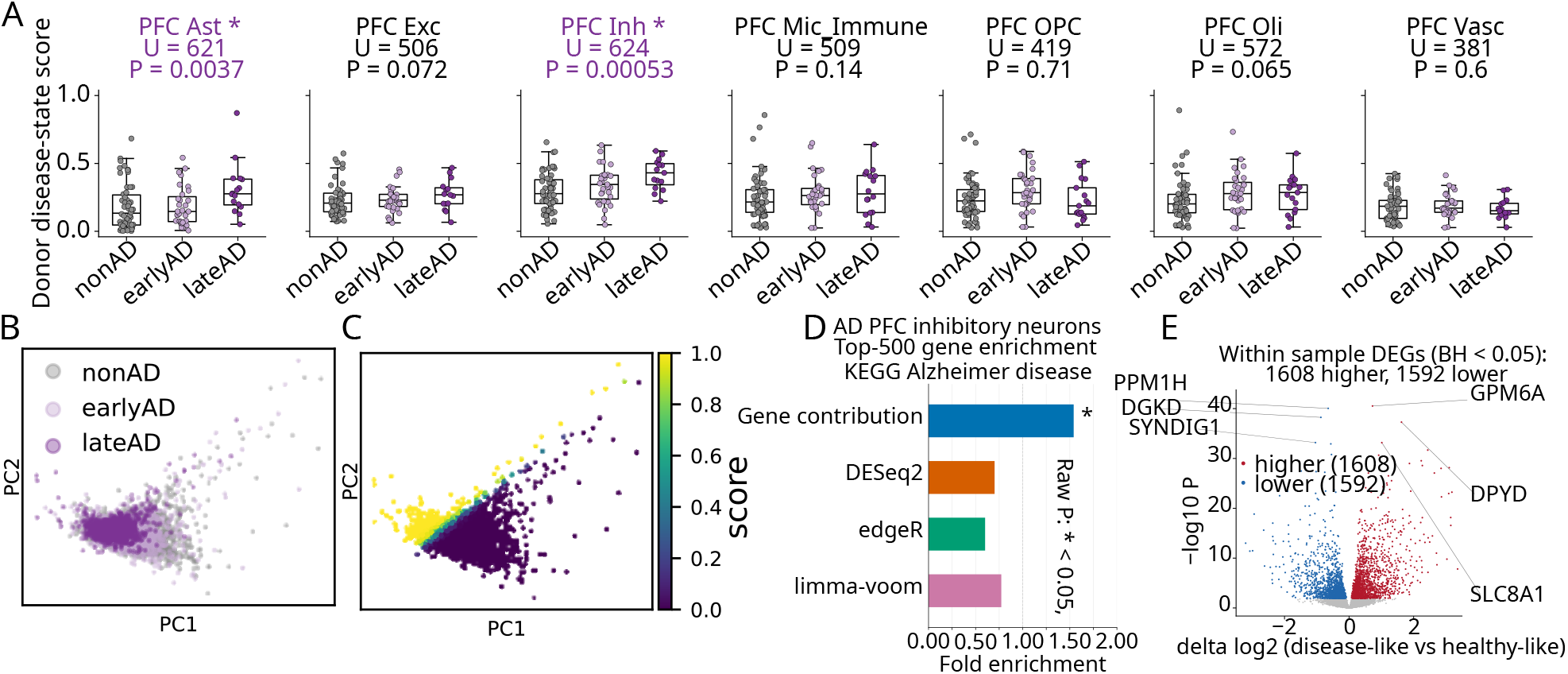
Disease-associated states in Alzheimer’s disease prefrontal cortex. (A) Donor scores across seven PFC cell classes, ordered nonAD, earlyAD and lateAD. nonAD and lateAD scores use leave-one-donor-out evaluation; earlyAD scores are averaged across frozen LODO models and shown descriptively. Points represent donors; boxes show medians and interquartile ranges, with whiskers extending to the most extreme observations within 1.5 interquartile ranges. *U* and raw *P* values are from exact one-sided Mann–Whitney tests of lateAD > nonAD; asterisks indicate nominal *P* < 0.05. (B,C) Principal-component views of the IDC embedding of PFC inhibitory neurons, colored by diagnostic group (B) or disease-like score (C); earlyAD nuclei were projected without fitting. (D) Over-representation of KEGG Alzheimer disease among the top 500 genes ranked by the IDC gene contribution score or the absolute case–control statistics defined in Fig. 4E. Bars show fold enrichment relative to each method’s ranked-gene background; the asterisk indicates raw hypergeometric *P* < 0.05. This analysis is unsigned and distinct from signed differential-expression GSEA. (E) Within-donor volcano plot across 15 lateAD donors, showing the reported Δ log_2_ effect-size summary versus − log_10_ *P*. Red and blue indicate genes higher (1,608) or lower (1,592) in disease-like cells at BH < 0.05; gray indicates other tested genes. Selected genes are labeled.

We next examined the cellular distribution of the AD-associated signal in inhibitory neurons. Although cells from non-AD, early-AD, and late-AD donors largely overlapped, non-AD cells occupied a broader region of the two-dimensional representation, including regions less populated by late-AD cells (Fig. 5B). Coloring cells by disease-like score revealed a gradient of disease-like states across this heterogeneous population (Fig. 5C). This pattern suggests that AD-associated changes may involve not only the emergence of disease-like states but also a reduced representation of cellular states prevalent in healthy donors.

To assess the biological relevance of the learned disease-associated direction, we examined enrichment of the KEGG Alzheimer disease pathway among the top 500 genes ranked by gene contribution scores. IDC showed significant enrichment (fold enrichment 1.54; hypergeometric *P* < 0.05), whereas the corresponding rankings from DESeq2, edgeR, and limma-voom showed substantially lower enrichment (Fig. 5D, Fig. <u>S7</u>). These results suggest that the direction learned by IDC captures transcriptional features associated with established AD biology.

We next identified genes associated with the inferred disease-like state using IDC’s within-donor differential expression analysis. Among the most significant genes in PFC inhibitory neurons, SYNDIG1, implicated in excitatory synapse development, showed lower expression in disease-like cells, whereas GPM6A, involved in neurite formation, and SLC8A1, encoding a sodium–calcium exchanger, showed higher expression (Fig. 5E). PPM1H, a Rab phosphatase, and DGKD, involved in lipid signaling, also showed lower expression. Together, these findings highlight transcriptional differences related to synaptic organization, calcium homeostasis, membrane trafficking, and lipid signaling along the inferred AD-associated cellular state.

## 4 Discussion

In case–control single-cell studies, disease often affects only some cells within a patient, while patients differ from controls in many ways besides disease. IDC uses the first fact to address the second. The mixture of affected and unaffected cells within a patient creates gene–gene covariance that healthy donors lack, and shifts that move all of a donor’s cells together leave this covariance unchanged. By learning from structure within donors, IDC scores cells, estimates each donor’s disease burden, and tests genes without batch labels.

IDC provides two views of disease at the gene level. Gene contribution scores show which genes define the learned disease axis. In beta cells and inhibitory neurons, the top-ranked genes were enriched for the KEGG type 2 diabetes and Alzheimer’s disease pathways, where case–control rankings showed weak or no enrichment. The within-donor test asks a different question: which genes change as cells become more disease-like along that axis. Because these genes need not define the axis themselves, this test captures a broader set of changes, including the metabolic, cell-adhesion and synaptic genes we observed. In simulations, it nearly doubled gene recovery relative to donor-level pseudobulk while keeping the false discovery rate close to the nominal level.

Working within donors also defines what IDC cannot see. An effect that shifts all of a patient’s cells equally cancels within the donor. This applies to batch, but also to biology, such as systemic hyperglycemia, circulating cytokines or medication. Case–control comparisons do capture such effects, but cannot separate them from batch and other donor-level differences. The two approaches therefore answer different questions. Between-donor analysis asks how patients differ from controls overall; IDC asks what distinguishes disease-like from healthy-like cells within the same tissue. Used together, they can separate patient-wide responses from changes confined to a subset of cells.

Several limitations remain. First, because donors are the units of replication, within-donor differential expression needs an adequate number of donors; in our benchmarks, this was at least ten per condition, although cell identification and burden estimation remained accurate in smaller cohorts. Second, technical variation among cells of the same donor, such as ambient RNA contamination or differences in nuclear quality, can still contribute to within-donor covariance. Excluding mitochondrial and ribosomal genes reduced this effect, and adding cell-level quality covariates to the covariance contrast is a natural extension. Third, IDC assumes that controls contain few affected cells; if control donors carry early pathology, as non-AD donors may, the contrast is weakened. Finally, we fitted IDC separately for each cell type. Linking disease programs across interacting cell types and across modalities is an important next step.

## Data and Code Availability

The type 2 diabetes data are single-nucleus multiome profiles of human pancreatic islets, available from the Gene Expression Omnibus under accession GSE200044 [19]. The Alzheimer’s disease data are ROSMAP single-nucleus RNA-seq profiles of prefrontal and entorhinal cortex from Liu et al. [20], available through the AD Knowledge Portal (Synapse) under the ROSMAP data use agreement (Synapse: syn52293442). Simulated data and the scripts that generated them are deposited with the code. IDC, the within-donor test, and all analysis scripts are available at https://github.com/cfg824157947/IDC-public (version v0.1.0), with environment files and a README.

## Acknowledgments

The authors are partially supported by National Institutes of Health grants R35GM150513.

## Supplementary Figures

**Figure S1:**
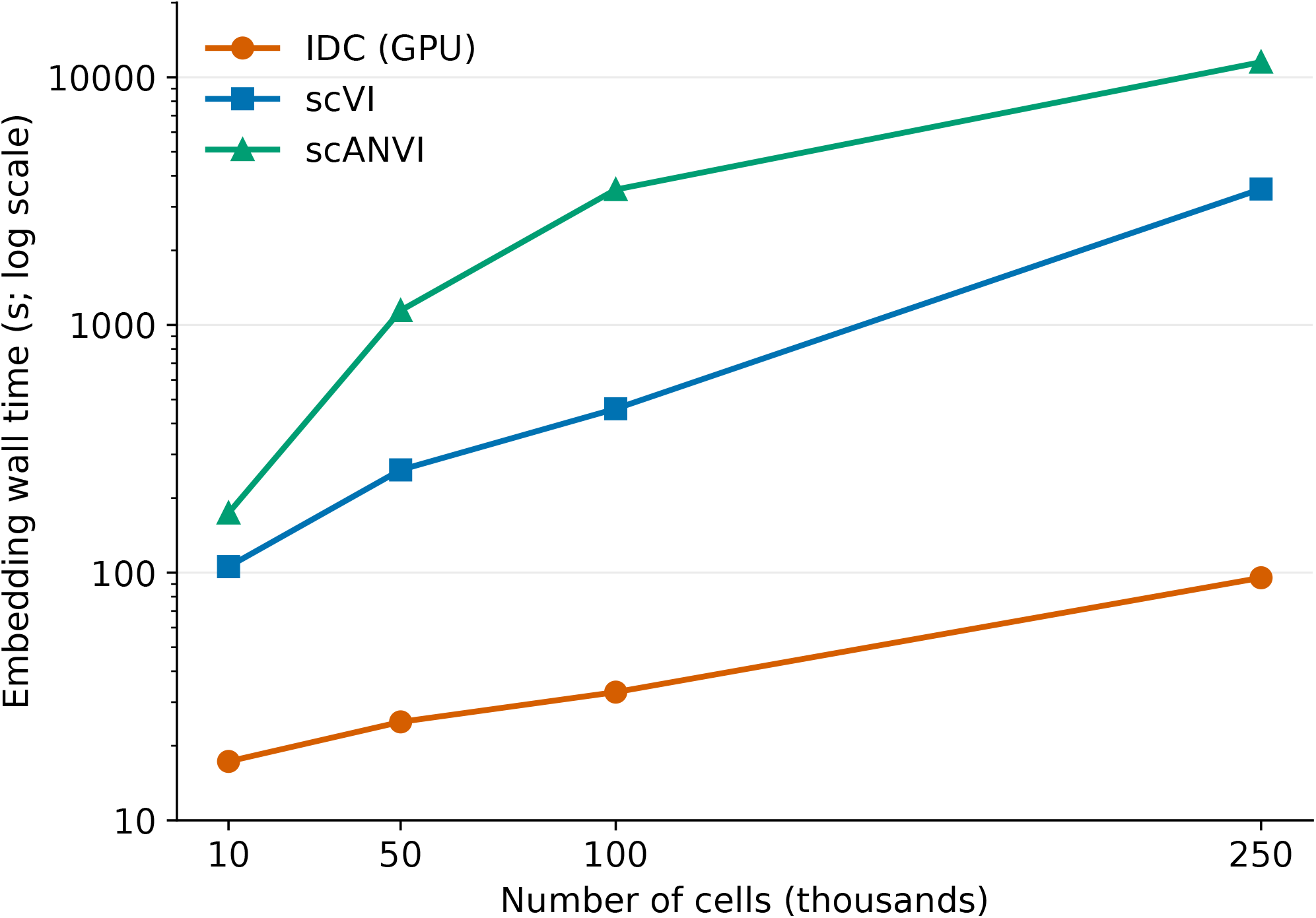
Embedding runtime across cell counts. Wall time for IDC (GPU), scVI, and scANVI representation generation using 10,000, 50,000, 100,000, and 250,000 cells with 18,000 genes on NVIDIA A100 80GB PCIe GPUs. The vertical axis is logarithmic. Points represent individual saved benchmark runs, with identical input checksums across methods at each cell count; lines connect measurements. Times use the saved `wall_seconds_from_started` field, measured after input loading through representation generation and output handling. PU learning and gene-level testing are excluded. No memory measurements are shown.

**Figure S2:**
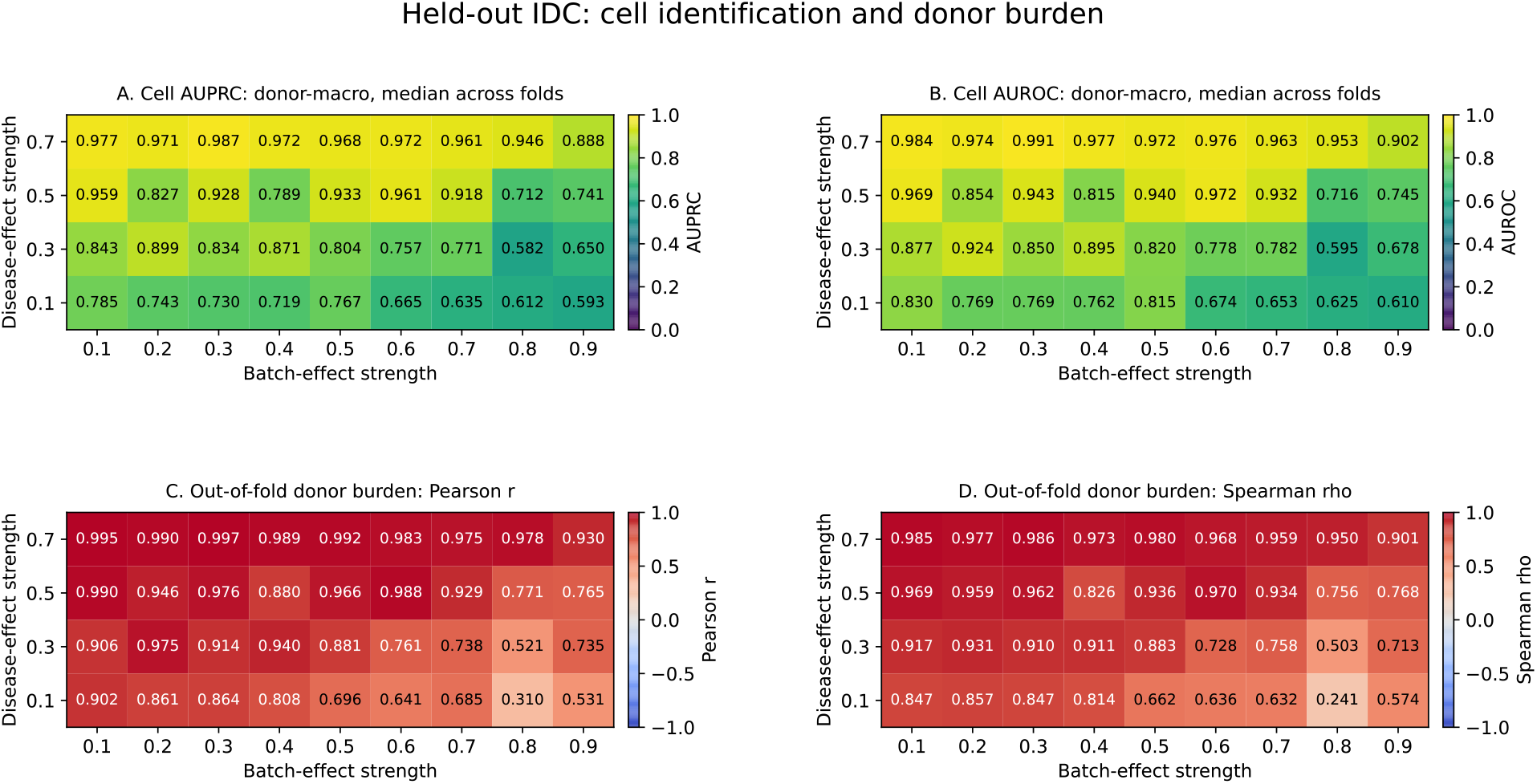
Held-out IDC validation across 36 simulation settings. Rows indicate disease-effect strength and columns batch-effect strength. (A,B) Cell-identification AUPRC and AUROC, calculated within disease donors and summarized by donor-macro performance and the median across five donor-disjoint folds. (C,D) Agreement between mean out-of-fold cell score and true affected-cell fraction across the 30 disease donors: Pearson correlation (C), Spearman correlation (D). Correlations use color limits of −1 to 1.

**Figure S3:**
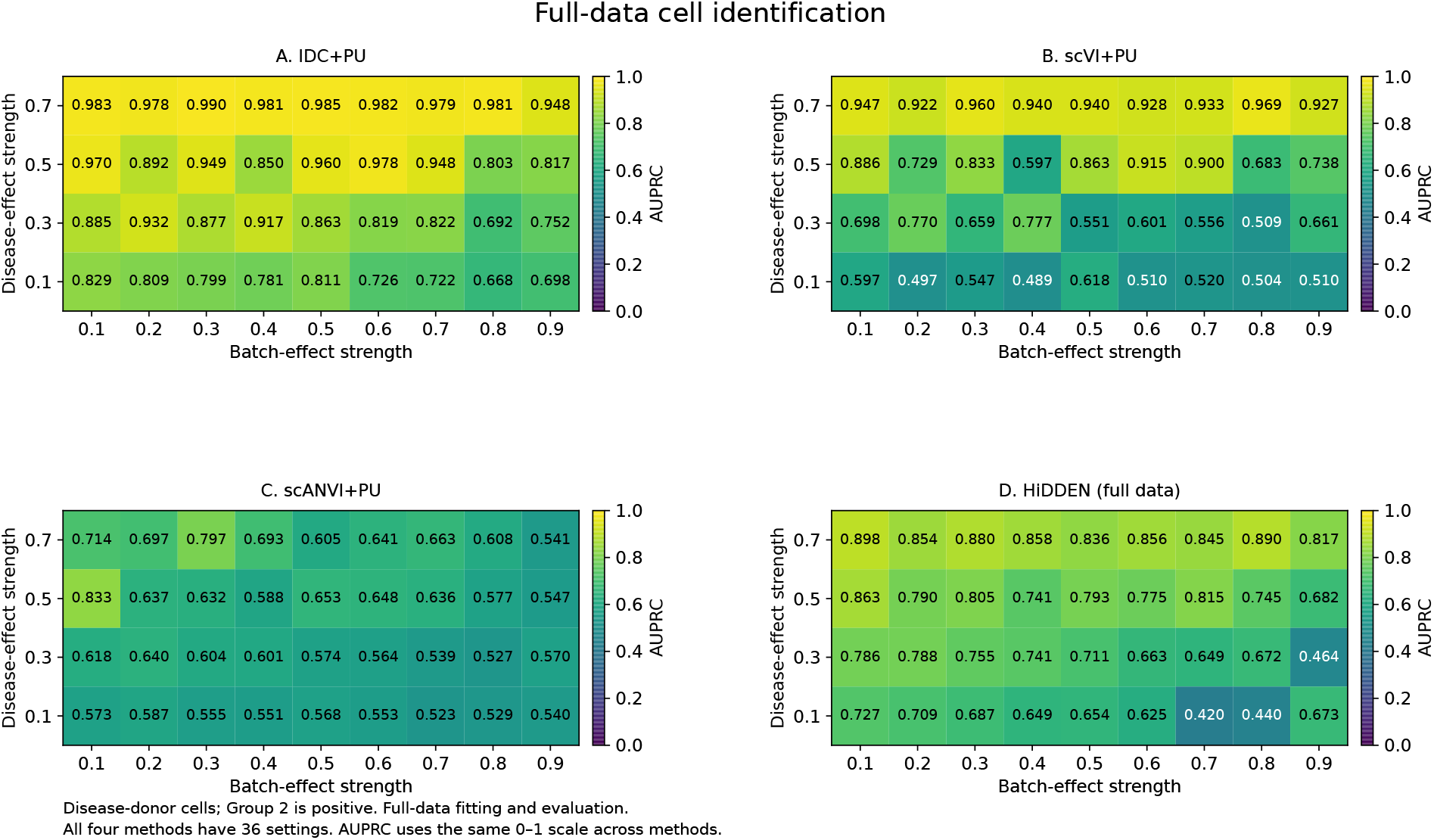
Full-data cell identification across 36 simulation settings. Disease-effect strengths form rows and batch-effect strengths form columns. Entries show AUPRC for identifying affected Group 2 cells among cells from disease donors using (A) IDC+PU, (B) scVI+PU, (C) scANVI+PU, and (D) HiDDEN. The same PU procedure is applied to the first three representations. All methods are fitted and evaluated on the full dataset; these are not held-out performance estimates. AUPRC color limits are 0–1 for all panels. Numerical annotations are rounded to three decimals; full-precision values are supplied in the accompanying table.

**Figure S4:**
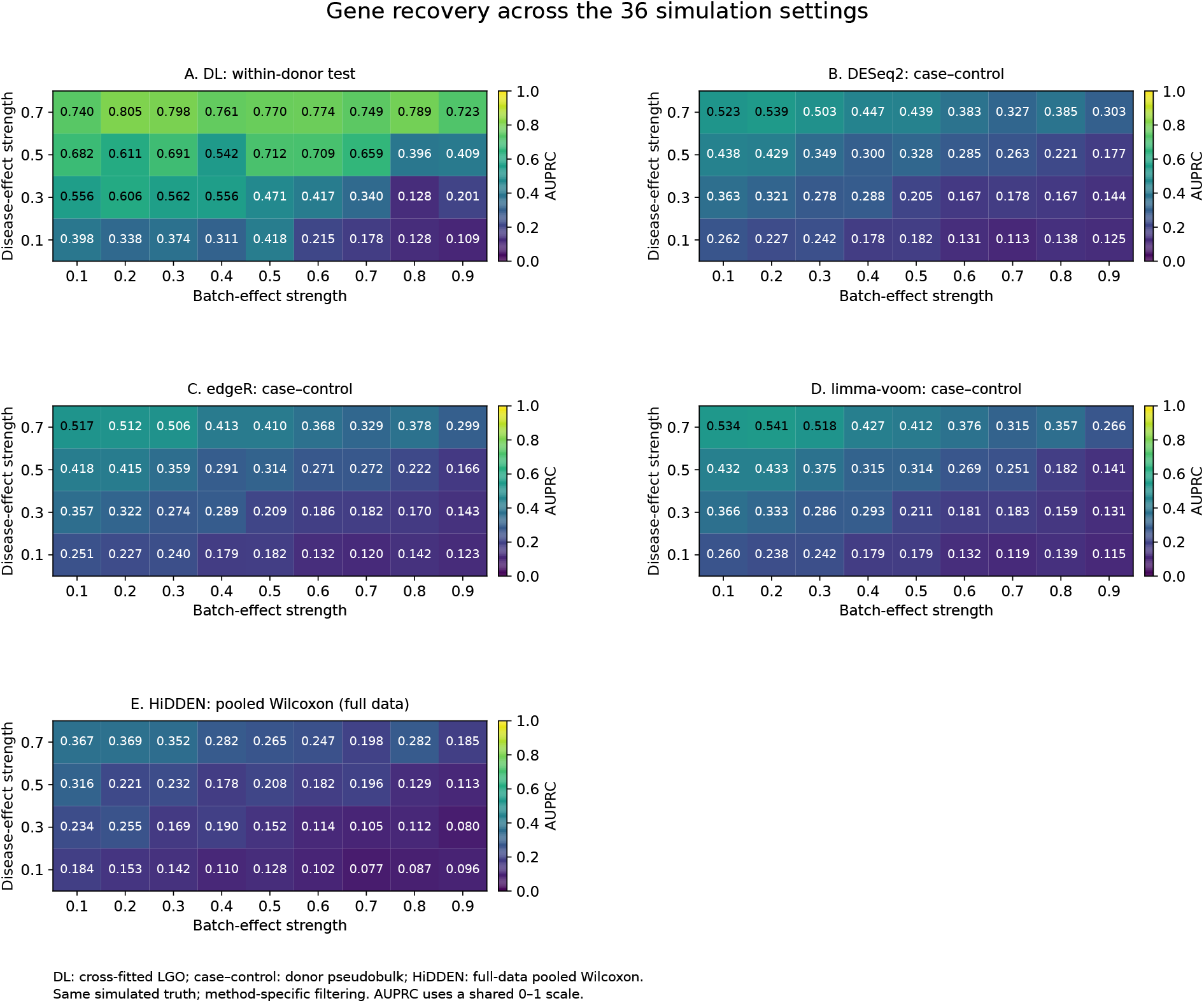
Gene-recovery AUPRC across 36 simulation settings. Panels show the IDC within-donor DerSimonian– Laird test (A), case–control donor-pseudobulk DESeq2 (B), edgeR (C), limma-voom (D), and HiDDEN pooled Wilcoxon testing of full-data refined labels (E). Rows indicate disease-effect strength and columns batch-effect strength. The within-donor test uses cross-fitted leave-gene-out scores and requires at least two fitted donors per tested gene. The existing all-gene AUPRC convention is preserved, including untested genes assigned *P* = 1 for the harmonized DL ranking. Methods use the same simulated gene truth but retain their established filtering and evaluation procedures. HiDDEN’s component count is selected by maximizing pooled BH-0.05 discoveries using its established full-data workflow; it is not a donor-held-out test. All panels use an AUPRC color scale of 0–1.

**Figure S5:**
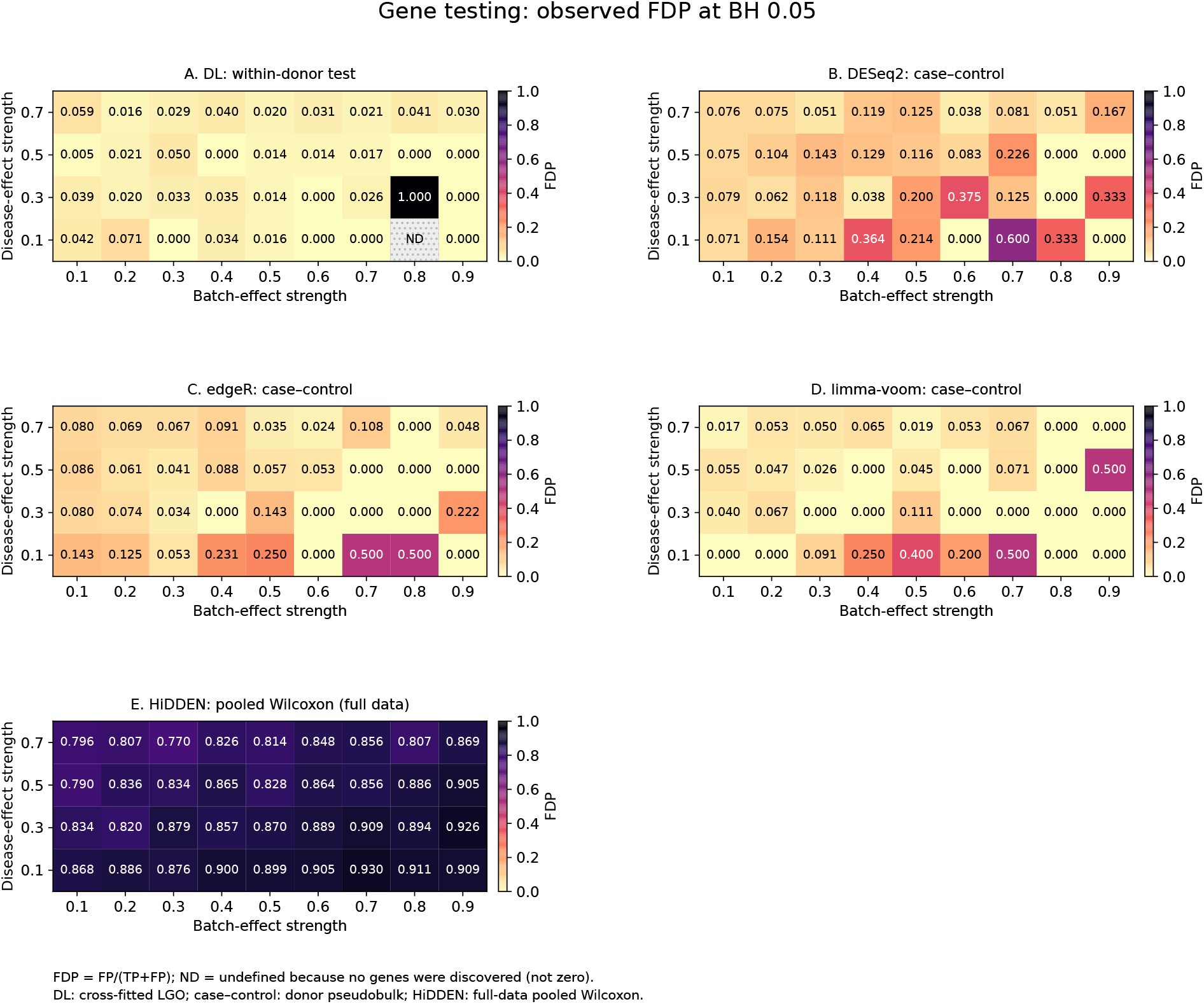
Observed false discovery proportion at BH 0.05 across 36 simulation settings. Panels show the IDC within-donor DerSimonian–Laird test (A), case–control DESeq2 (B), edgeR (C), limma-voom (D), and HiDDEN pooled full-data Wilcoxon test (E), with the same protocols as the gene-recovery figure. Rows indicate disease-effect strength and columns batch-effect strength. Observed FDP is FP /(TP + FP) when at least one gene is discovered; ND indicates no discoveries and undefined FDP, not zero. All panels share a 0–1 scale. For DL, 33 of the 35 settings with discoveries had observed FDP at most 0.06, with a median of 0.020. One setting yielded no discoveries. At disease-effect strength 0.3 and batch-effect strength 0.8, IDC identified one false-positive gene, resulting in an FDP of 1.0. Numerical annotations are rounded to three decimals, and full-precision values are provided separately.

**Figure S6:**
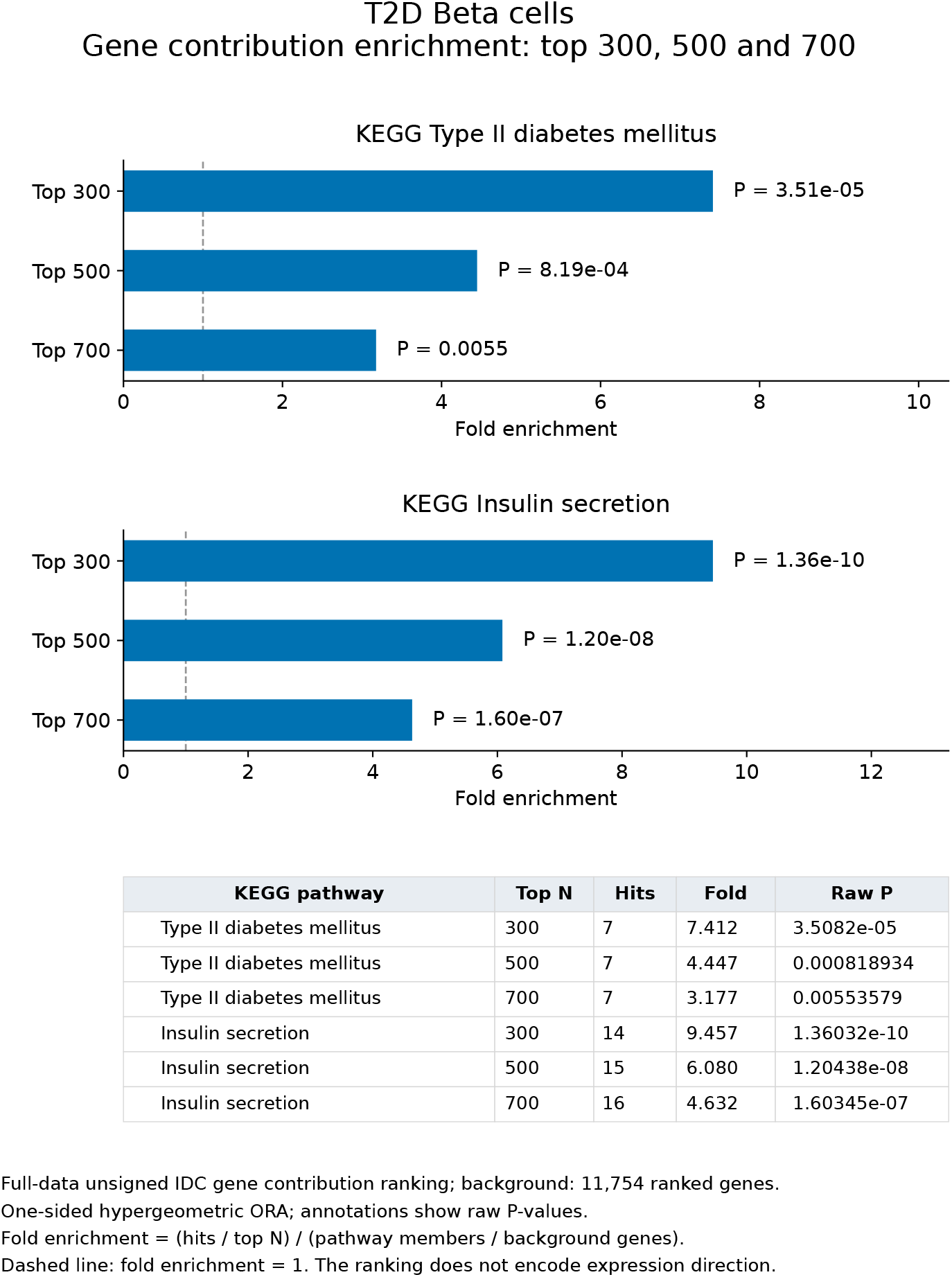
KEGG pathway enrichment among genes contributing to the T2D-associated cell state. The top 300, 500, and 700 genes ranked by unsigned disease-state gene contribution scores from the full-data IDC fit in pancreatic beta cells were tested for overrepresentation of the KEGG Type II diabetes mellitus and Insulin secretion pathways. Bars show fold enrichment relative to all genes in the contribution ranking; the dashed line indicates no enrichment (fold enrichment = 1). Annotations report raw, one-sided hypergeometric P-values. The table lists pathway overlaps and enrichment statistics. Gene contribution scores characterize the learned disease direction and do not indicate whether expression is higher or lower in disease-like cells.

**Figure S7:**
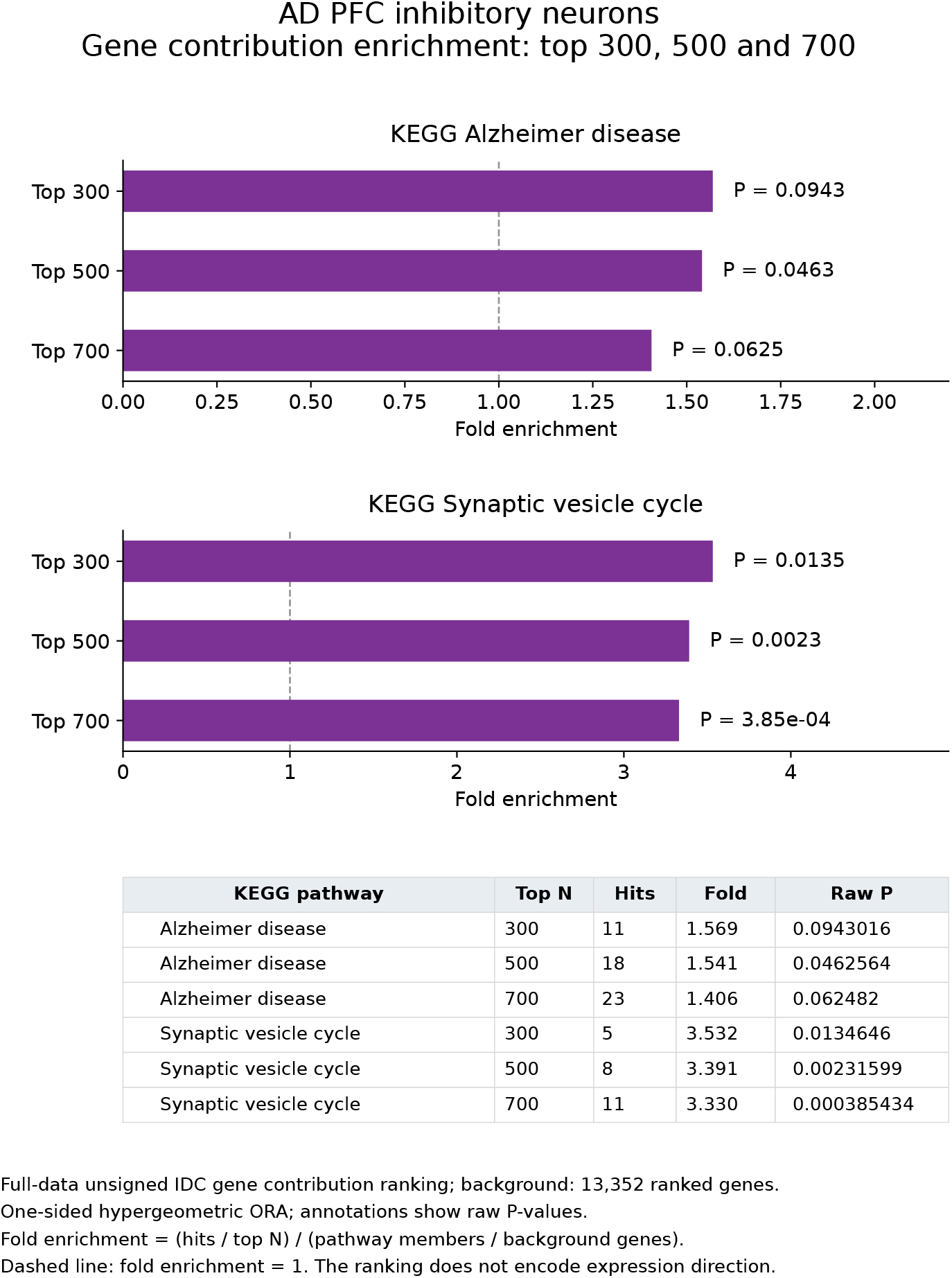
KEGG pathway enrichment among genes contributing to the AD-associated cell state. The top 300, 500, and 700 genes ranked by unsigned disease-state gene contribution scores from the full-data IDC fit in PFC inhibitory neurons were tested for overrepresentation of the KEGG Alzheimer disease and Synaptic vesicle cycle pathways. Bars show fold enrichment relative to all genes in the contribution ranking; the dashed line indicates no enrichment (fold enrichment = 1). Annotations report raw, one-sided hypergeometric P-values. The table lists pathway overlaps and enrichment statistics. Gene contribution scores characterize the learned disease direction and do not indicate whether expression is higher or lower in disease-like cells.

